# *In vivo* dissection of the mouse tyrosine catabolic pathway with CRISPR-Cas9 identifies modifier genes affecting hereditary tyrosinemia type 1

**DOI:** 10.1101/2023.09.29.559947

**Authors:** Jean-François Rivest, Sophie Carter, Claudia Goupil, Denis Cyr, Roth-Visal Ung, Dorothée Dal Soglio, Fabrice Mac-Way, Paula J. Waters, Massimiliano Paganelli, Yannick Doyon

**Author notes:** Address correspondence to: Yannick Doyon, Ph.D. Centre de recherche du CHU de Québec – Université Laval 2705, boulevard Laurier, T-3-67 Québec, QC G1V 4G2 CANADA.

## Abstract

Hereditary tyrosinemia type 1 is an autosomal recessive disorder caused by mutations (pathogenic variants) in fumarylacetoacetate hydrolase, an enzyme involved in tyrosine degradation. Its loss results in the accumulation of toxic metabolites that mainly affect the liver and kidneys and can lead to severe liver disease and liver cancer. Tyrosinemia type 1 has a global prevalence of approximately 1 in 100,000 births but can reach up to 1 in 1,500 births in some regions of Québec, Canada. Mutating functionally related ‘modifier’ genes (*i.e*., genes that, when mutated, affect the phenotypic impacts of mutations in other genes) is an emerging strategy for treating human genetic diseases. *In vivo* somatic genome editing in animal models of these diseases is a powerful means to identify modifier genes and fuel treatment development. In this study, we demonstrate that mutating additional enzymes in the tyrosine catabolic pathway through liver-specific genome editing can relieve or worsen the phenotypic severity of a murine model of tyrosinemia type 1. Neonatal gene delivery using recombinant adeno-associated viral vectors expressing *Staphylococcus aureus* Cas9 under the control of a liver-specific promoter led to efficient gene disruption and metabolic rewiring of the pathway, with systemic effects that were distinct from the phenotypes observed in whole-body knockout models. Our work illustrates the value of using *in vivo* genome editing in model organisms to study the direct effects of combining pathological mutations with modifier gene mutations in isogenic settings.

## INTRODUCTION

Inborn errors of metabolism (IEMs) are inherited genetic disorders caused by disruptions in a specific enzymatic reaction within a metabolic pathway that induce pathology through toxic metabolite accumulation or deficiencies in downstream metabolites^1^. First described by Archibald Garrod over 100 years ago, IEMs are frequently considered to be monogenic diseases^2^. However, this classification may over-simplify the biological reality, as patients often present a spectrum of phenotypes^3–5^. Importantly, modifier genes can profoundly influence the phenotype associated with mutation(s) (pathogenic variants) at a primary “disease-causing” gene locus^3–8^. Genome editing therapies targeting modifier genes are already showing great promise in clinical trials, particularly for hemoglobinopathies^9–11^. Liver-specific base editing of *PCSK9* is also currently being investigated in a clinical trial to treat familial hypercholesterolemia^12, 13^. Thus, identifying and characterizing modifier genes can provide valuable mechanistic information and spark the development of novel therapeutics.

The phenylalanine and tyrosine degradation pathway is notable historically as it is linked to the description of the first inborn error of metabolism, alkaptonuria^14^. Loss-of-function of each enzyme in this pathway leads in a different IEM (**Figure 1A**). In the first step, phenylalanine is converted to tyrosine by phenylalanine hydroxylase (PAH). Loss of PAH activity leads to phenylketonuria; a disease characterized by intellectual disability and seizures (OMIM 261600). Tyrosine is then converted to 4-hydroxyphenylpyruvate (4- HPP) by tyrosine aminotransferase (TAT). TAT inactivation results in tyrosinemia type II, which causes painful corneal lesions, skin disease, and intellectual disability (OMIM 276600). In the next step, 4-HPP is converted to homogentisic acid (HGA) by 4- hydroxyphenylpyruvate dioxygenase (HPD). Its loss-of-function results in tyrosinemia type III, a disease characterized by intellectual disability, seizures, and intermittent ataxia (OMIM 276710). In the fourth reaction, HGA is converted to maleylacetoacetate (MAA) by homogentisate 1,2-dioxygenase (HGD). Patients with inactive HGD display high HGA levels and develop alkaptonuria, the prototypical IEM described by Garrod^2^, which results in arthritis, heart valve and kidney issues, and pigmentation changes to the cartilage and urine (OMIM 203500). In the penultimate step in tyrosine degradation, MAA is converted to fumarylacetoacetate (FAA) by glutathione S-transferase zeta 1 (GSTZ1), also known as maleylacetoacetate isomerase (MAAI). Individuals with GSTZ1 deficiency display mild hypersuccinylacetonemia which appears to be clinically insignificant^15, 16^ (OMIM 617596). Finally, fumarylacetoacetate hydrolase (FAH) converts FAA into fumarate and acetoacetate, which are used for energy production by the tricarboxylic acid cycle and reconverted to acetyl-CoA respectively.

**Figure 1.**
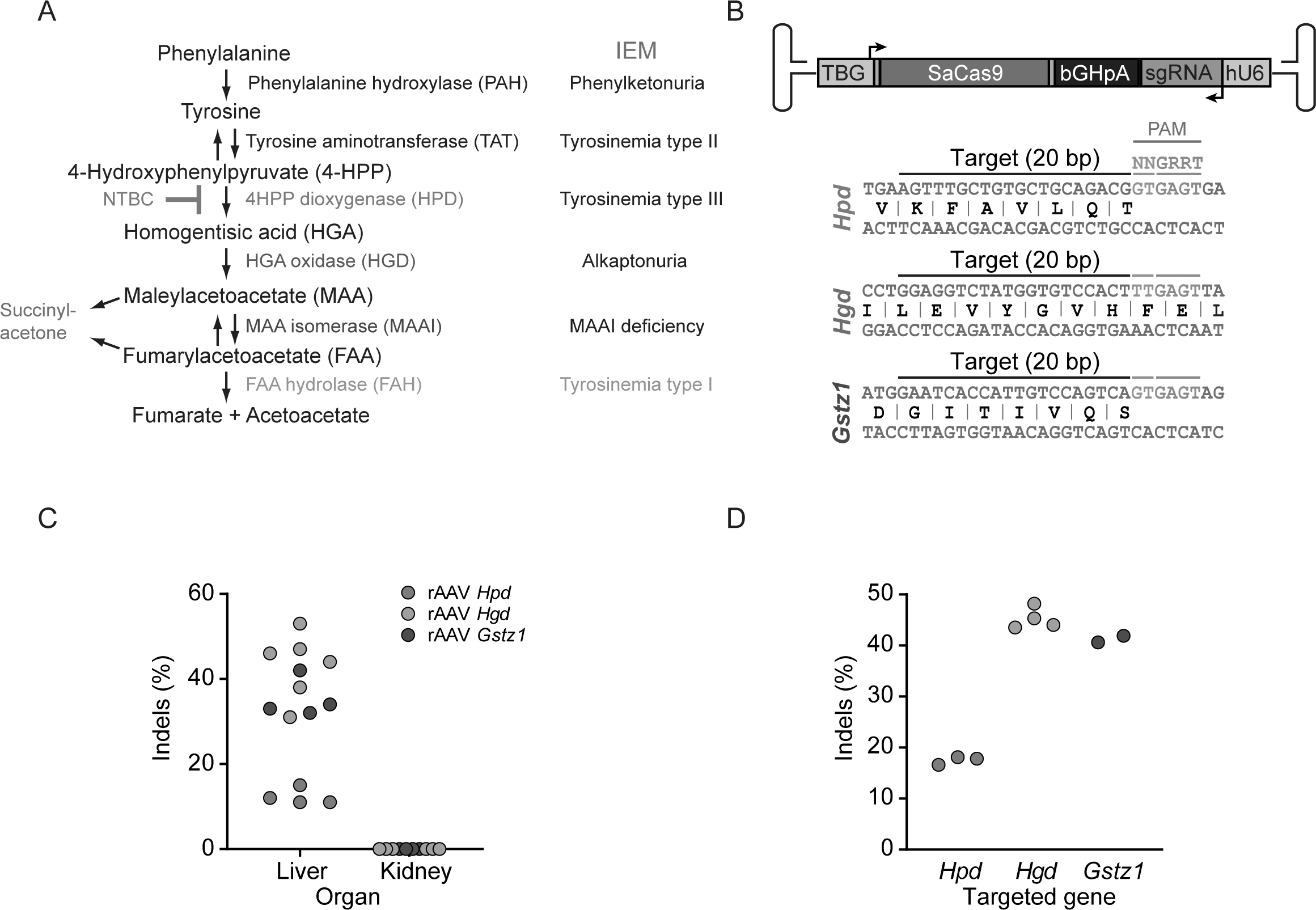
*In vivo* genome editing of the tyrosine catabolic pathway by rAAV8-SaCas9. (A) The tyrosine degradation pathway and associated inborn errors of metabolism (IEMs). The catabolic reaction inhibited by NTBC is indicated in red. 4-HPP, 4-hydroxyphenylpyruvate; HGA, homogentisic acid; MAA, maleylacetoacetate; FAA, fumarylacetoacetate. (B) **Top:** Schematic of the rAAV-SaCas9 vector. The thyroxine-binding globulin (TBG), bovine growth hormone polyadenylation (bGHpA), and hU6 promoter sequences are indicated. **Bottom:** Sequences of the sgRNAs targeting *Hgd, Hgd* and *Gstz1* chosen for *in vivo* studies. The protospacer adjacent motifs (PAMs) are annotated. The two blue nucleotides in the *Gstz1* target site are mismatches compared to the human sequence. (C) Neonatal C57/Bl6 mice were injected with 5×10^10^ vector genomes (VGs) of rAAV8-SaCas9 into the retro-orbital sinus, weaned at 21 days old, and sacrificed at the following age: *Hpd,* 40 days; *Hgd,* 40 days; *Gstz1*, 77 days. Genomic whole-liver (*Hpd*, *Hgd*, and *Gstz1*) and kidney (*Hgd* and *Gstz1*) DNA was extracted, and Surveyor assays were used to determine indel frequencies. Each symbol represents a different animal. A mouse injected with saline was used as the negative control for the Surveyor assay. (D) Neonatal male *Fah^-/-^* mice were injected with 5×10^10^ VGs of rAAV8-SaCas9 into the retro-orbital sinus, weaned at 21 days old, and sacrificed at 28 days old after continuous NTBC treatment. TIDE assays were used to determine the frequency of indels. A mouse injected with saline was used as the negative control for the TIDE assay.

Among IEMs affecting this pathway, one of the most severe is undoubtedly hereditary tyrosinemia type 1 (HT-I), a disorder caused by the absence of a functional *FAH* allele (OMIM 276700)^17^. Loss of FAH activity can lead to hepatic failure with renal and neurological comorbidities, along with a high risk of developing hepatocellular carcinoma^18, 19^. HT-I results from the accumulation of tyrosine and its toxic metabolites, such as FAA, MAA, and succinylacetone (SA), a by-product of both FAA and MAA degradation and the diagnostic marker for HT-I^20^. However, the molecular mechanisms by which these compounds damage the liver and kidneys are poorly characterized^18, 20^. HT-I is progressive and life-threatening if left untreated. Standard care includes diet therapy to limit phenylalanine and tyrosine intake and lifelong treatment with 2-(2-nitro-4- trifluoromethylbenzoyl)-1,3-cyclohexanedione (NTBC; also known as nitisinone), a potent inhibitor of the upstream enzyme HPD^21^, to prevent toxic metabolite accumulation in the liver and kidneys^22–24^ (**Figure 1A**). Importantly, non-compliance with NTBC and diet treatment is a serious challenge for the clinical management of HT-I and results in higher risks of patients developing hepatocellular carcinoma, as well as painful corneal lesions and neurological crises due to high circulating tyrosine levels^25^. While this catabolic pathway is well characterized at the molecular genetic level, the broad spectrum of phenotypes and disease severity observed in patients with HT-I—even within families—is not completely understood^26^.

Several knockout mouse models have been created to evaluate the interplay between the different genes in the phenylalanine and tyrosine degradation pathway and their impacts on various disease phenotypes^27–31^. The *Fah*^Δexon5^ (*Fah*^-/-^) mouse is a well-established murine model for HT-I pathophysiology^23, 27^. In these mice, neonatal lethality can be prevented by early treatment with NTBC, which completely corrects the metabolic liver disease and results in a phenotype analogous to human tyrosinemia type I^23^. Upon NTBC withdrawal, however, disease progression resumes leading to death in a short period of time^32^. This experimental system offers an opportunity to test treatment options *in vivo*. Moreover, in both mice and humans, transplanted or genetically corrected hepatocytes have a potent selective growth advantage in the mutant liver lowering the initial threshold for treatment efficacy ^27, 32–36^. Hence, this model has become a staple for hepatic gene therapy research^32^. Whole*-*body double knockout mouse studies have revealed that full metabolic blocks at different points upstream in the pathway greatly affect the disease phenotype. For example, while *Fah*^-/-^ *Hpd*^-/-^ and *Fah*^-/-^ *Hgd*^-/-^ animals survived without NTBC, *Fah^-/-^ Gstz1^-/-^* mice displayed an even more severe phenotype than *Fah^-/-^* mice^30, 37, 38^ (**Figure 1A**). Accordingly, both short hairpin RNA (shRNA)-mediated knockdown and CRISPR-Cas9- directed inactivation of *Hpd* in the liver of *Fah^-/-^* animals enabled the survival after NTBC withdrawal, suggesting that metabolically blocking the pathway in the liver is sufficient for a systemic therapeutic effect^39–42^. Still, the impact of liver-specific inactivation of *Hgd* and *Gstz1* via *in vivo* genome editing in the HT-I mouse model has yet to be described. To test this, we used rAAV8-mediated delivery of *Streptococcus aureus* Cas9 (SaCas9)^43^ to inactivate *Hpd*, *Hgd*, and *Gstz1* in neonatal *Fah*^Δexon5^ (*Fah*^-/-^) mouse (**Figure 1A**). We did not target *Tat*, as its loss produces a more severe phenotype in mice than the loss of *Hpd*, *Hgd*, and *Gstz1*, according to phenotyping data from the the International Mouse Phenotyping Consortium^27, 30, 31, 38, 44^. We found that, while targeting *Hpd* and *Gstz1* yielded outcomes similar to those observed in whole-body double mutant mice, targeting *Hgd* led to systemic and lethal effects in contrast to the genetic suppression observed in the conventional *Fah^-/-^ Hgd^-/-^* model. This work highlights the value of somatic genome editing in animals for modelling human disorders^45^.

## RESULTS

### Potent *in vivo* editing in the liver of neonatal mice via AAV8-mediated delivery of SaCas9

Since most of the enzymes in the tyrosine catabolic pathway are expressed mainly in the liver and kidneys, we hypothesized that liver-specific gene disruption would produce a systemic impact on metabolic functions. This can be achieved in neonatal mice using CRISPR-Cas9 delivered via rAAV8 injected systemically to target hepatocytes^46^. In this system, the expression of a single guide RNA (sgRNA) targeting a specific gene is driven by the ubiquitous U6 promoter, and SaCas9 is expressed from a liver-specific thyroxine-binding globulin (TBG) promoter^43^ (**Figure 1B**). Following target cleavage in mouse hepatocytes, DNA repair mainly occurs through the error-prone non-homologous end joining pathway (NHEJ), creating insertions and deletions (indels) that disrupts the target gene^47^.

Using CRISPOR^48^, we designed several SaCas9 sgRNAs against *Hpd, Hgd,* and *Gstz1* (the gene encoding MAAI, **Figure 1A**) (**Table S1)**. We selected sgRNAs that targeted essential protein domains and had few predicted off-target effects on the mouse genome to increase the likelihood of generating inactivating mutations^49–51^. We identified highly active sgRNAs for all three targets by transient transfection in mouse neuroblastoma cells (**Figure S1)**. Nucleases with perfectly conserved target sites and protospacer adjacent motif (PAM) sequences between the mouse and human genomes were chosen for *Hpd* and *Hgd* but the *Gstz1* target site differs by 2bp **(Figure 1B)**. All-in-one rAAV vectors expressing both an sgRNA targeting *Hpd*, *Hgd,* or *Gstz1* and hepatocyte-specific SaCas9 (*via* the TBG promoter) were constructed and produced as hepatotropic rAAV serotype 8 (rAAV8) vectors **(Figure 1B)**^43, 52–54^. Next, neonatal (2-day-old) male C57BL/6N mice were injected into the retro-orbital sinuses with 5×10^10^ vector genomes (VGs) of rAAV8 targeting *Hpd, Hgd,* or *Gstz1*, then sacrificed 40 (*Hpd* and *Hgd*) or 77 (*Gstz1*) days later. Genomic whole-liver (*Hpd, Hgd,* and *Gstz1*) and kidney (*Hgd* and *Gstz1*) DNA was extracted, and Surveyor assays were used to determine indel frequencies^55^. Gene disruption was robust from mouse to mouse and generally efficient, ranging from 11–47% depending on the target in the liver (**Figure 1C)**. Since genomic DNA was extracted from whole livers and gene editing was limited to the hepatocytes (which comprise ∼70% of the liver’s mass^56^), the disruption efficiency is likely an underestimate. No editing was detected in the kidneys as expected when using rAAV8 and a liver-specific promoter to drive SaCas9 expression (**Figure 1C)**. We then repeated this experiment in 2-day-old male *Fah^-/-^* mice maintained on NTBC until sacrifice at 28 days old and observed similar levels of gene disruption as determined by the tracking of indels by decomposition (TIDE) assay^57^ (**Figure 1D)**. The indel profiles obtained with the TIDE assay indicated the presence of out-of-frame mutations likely to result in gene inactivation (**Figure S1).** These editing rates are similar to previous *in vivo* editing studies indicating that our system is fully functional in this context^43, 58^.

### *In vivo* inactivation of various enzymes in the tyrosine catabolic pathway differentially impacts metabolic outcomes and survival in HT-I mice

The impacts of liver-specific *in vivo* genome editing of *Hpd, Hgd,* and *Gstz1* (the gene encoding MAAI, **Figure 1A**) in HT-I mice were then determined upon NTBC withdrawal. Neonatal male *Fah^-/-^*mice treated with NTBC were injected at 2 days old with rAAV8- SaCas9 targeting *Hpd, Hgd,* or *Gstz1* (or saline as a control), and NTBC was withdrawn between 4–7 weeks old (**Figure 2A**). As expected, all mice treated with the nuclease targeting *Hpd* displayed normal lifespans, weights, and glycemia (blood glucose levels) post-NTBC removal, while saline-treated animals were sacrificed ∼3 weeks after NTBC removal, when meeting the weight loss criterion (**Figure 2B–D**). Glycemia and weight gains were normalized for over 14 weeks after NTBC withdrawal in *Hpd*-treated mice and they were kept alive without NTBC for 1 year (**Figure S2, Table S2**). In sharp contrast, *Fah^-/-^* mice treated with rAAV8-SaCas9 targeting either *Hgd* or *Gstz1* experienced sudden weight loss and hypoglycemia and died approximately 3 and 5 days post-NTBC removal, respectively (**Figure 2B–D**). We measured urine succinylacetone (SA) and homogentisic acid (HGA) levels respectively 24 hours before NTBC removal and 34 days after NTBC removal (*Hpd*) or the predicted time of death (*Hgd* and *Gstz1*). Before NTBC removal, the urine SA levels of the different treatment groups were broadly similar (**Figure 2E, Table S3**). The SA levels of mice treated with the *Hpd*-targeting vector decreased post-NTBC removal to only slightly above the detection limit indicating that the phenotype was normalized (**Figure 2E, Table S3**). Notably, mice treated with the *Gstz1*-targeting vector showed marked increases in urine SA after drug withdrawal that were equal to or higher than saline treated controls (**Figure 2E, Table S3).** Urine HGA levels were undetectable in all animals except in mice treated with the vector targeting *Hgd*, which experienced a considerable increase following NTBC withdrawal (**Figure 2F, Table S4**). These biochemical changes correspond to the step blocked in the pathway (**Figure 1A**). TIDE assays on the livers of sacrificed mice treated with the vectors targeting *Hgd* or *Gstz1* showed editing levels comparable to those seen in our previous tests with healthy animals (**Figures 2G and 1C,D**). Collectively, these data indicate that somatic genome editing can rewire the liver metabolism in HT-I mice to either suppress or enhance the disease.

**Figure 2.**
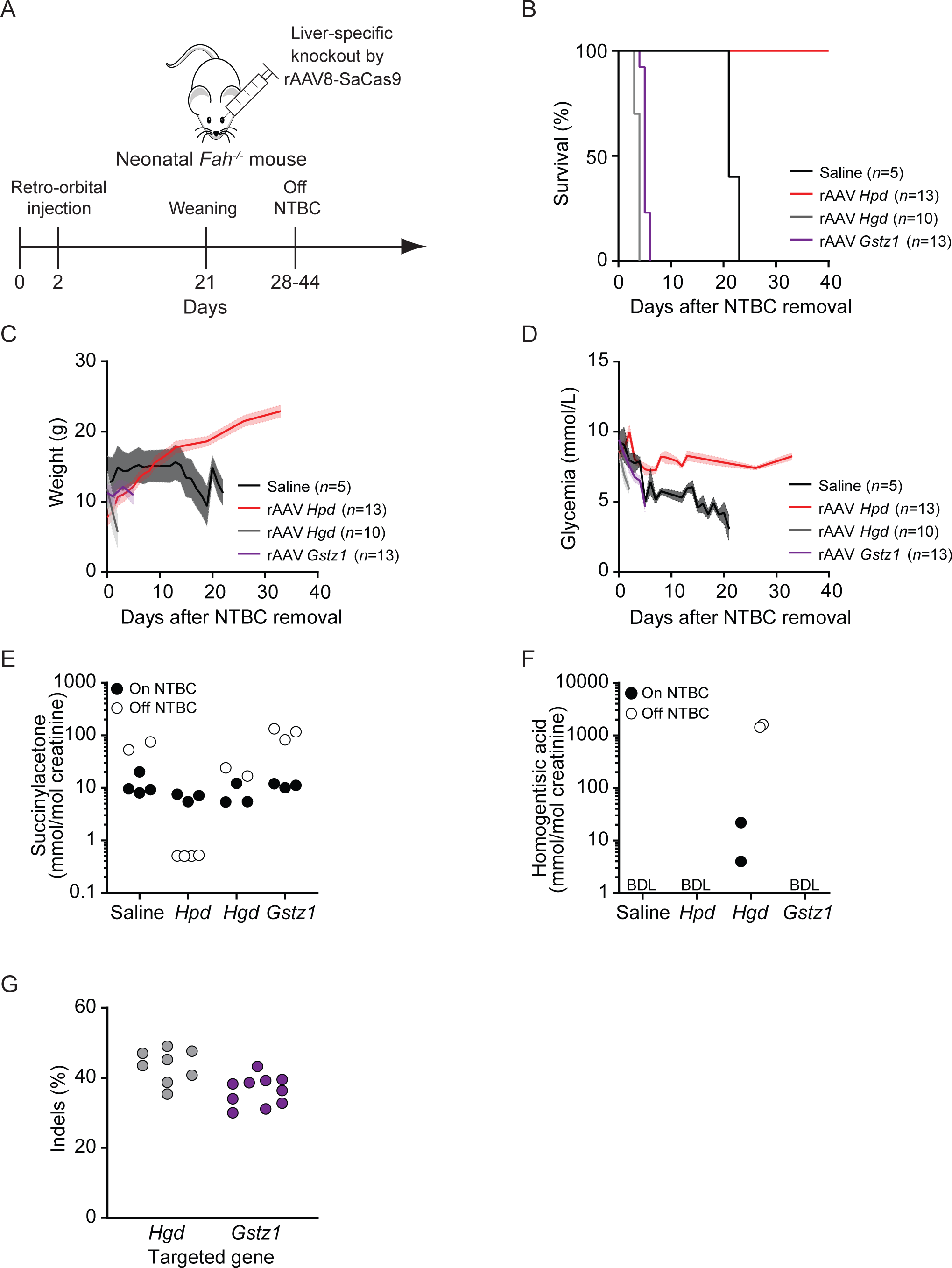
Metabolic rewiring through liver-specific knockout of *Hpd, Hgd,* and *Gstz1* in *Fah^-^*/- mice. (A) Experimental design for *in vivo* editing. Neonatal (2-day-old) male *Fah^-/-^* mice were injected with 5×10^10^ VGs of rAAV8-SaCas9 or saline into the retro-orbital sinus and weaned at 21 days old. NTBC treatment was stopped at different time points; 28 days (saline, *n*=3; *Hpd, n*=7; *Hgd*, *n*=3; *Gstz1*, *n*=3), 30 days (*Hpd, n*=6; *Gstz*, *n*=3), 41 days (saline, *n*=2; *Gstz1*, *n*=7), or 44 days (*Hgd*, *n*=7). Since the period elapsed between weaning and NTBC removal did not affect the outcome, mice from each treatment group were combined in the graphs shown in (B-D). The numbers of mice per group (*n*) and rAAV targets are indicated. (B) Survival following NTBC removal. Mice were sacrificed after losing 20% of their body weight. (C) Body weight was measured daily following NTBC removal. Solid lines indicate the mean and shaded areas denote the standard error of the mean (SEM). (D) Glycemia was monitored in non-fasted mice. (E–F) Urine succinylacetone (E) and homogentisic acid (F) levels were determined 24 hours before NTBC removal and again 24 hours before the predicted time of sacrifice (1, 4, and 19 days post- NTBC removal for mice treated with *Hgd* rAAV8, *Gstz1* rAAV8, and saline, respectively. Levels were determined in *Hpd* rAAV8-treated mice 34 days after NTBC removal. Samples were collected over 24 hours using metabolic cages containing 2-5 mice (**Tables S3 and S4**). The detection limits for succinylacetone and homogentisic acid were 0.1 mmol/mol and approximately 3 mmol/mol creatinine, respectively. BDL: below detection limit. (G) Genomic DNA was extracted from whole livers of mice treated with *Hgd-* and *Gstz1-*targeting vectors meeting the sacrifice endpoint, and Surveyor assays were used to determine indel frequencies. Each symbol represents a different animal. A mouse injected with saline was used as the negative control for the Surveyor assay.

As kidneys of HT-I mice are sensitive to cytotoxicity^23, 27, 30, 59^, we performed SDS-PAGE analysis of urine samples and observed a massive increase in serum albumin (also known as albuminuria) in the animals treated with the vectors targeting *Hgd* and *Gstz1* following NTBC removal, indicative of kidney disease^60^ (**Figure 3C**). At harvest, the kidneys of mice treated with the *Hgd*-targeting vector were pale, enlarged, and displayed severe diffuse tubular damage with necrosis, which is consistent with HGA-induced toxic tubular cell injury leading to acute renal insufficiency (**Figures 3A, top panel** and **3B**). Those treated with the vector targeting *Gstz1* had slight tubular damage mainly in the cortical area, similarly to saline-treated controls (**Figure 3A, top panel**). A group of *Hpd*-targeted mice sacrificed 35 days after NTBC withdrawal displayed normal tubules and glomeruli, suggesting preserved kidney function (**Figure 3A, top panel**).

**Figure 3.**
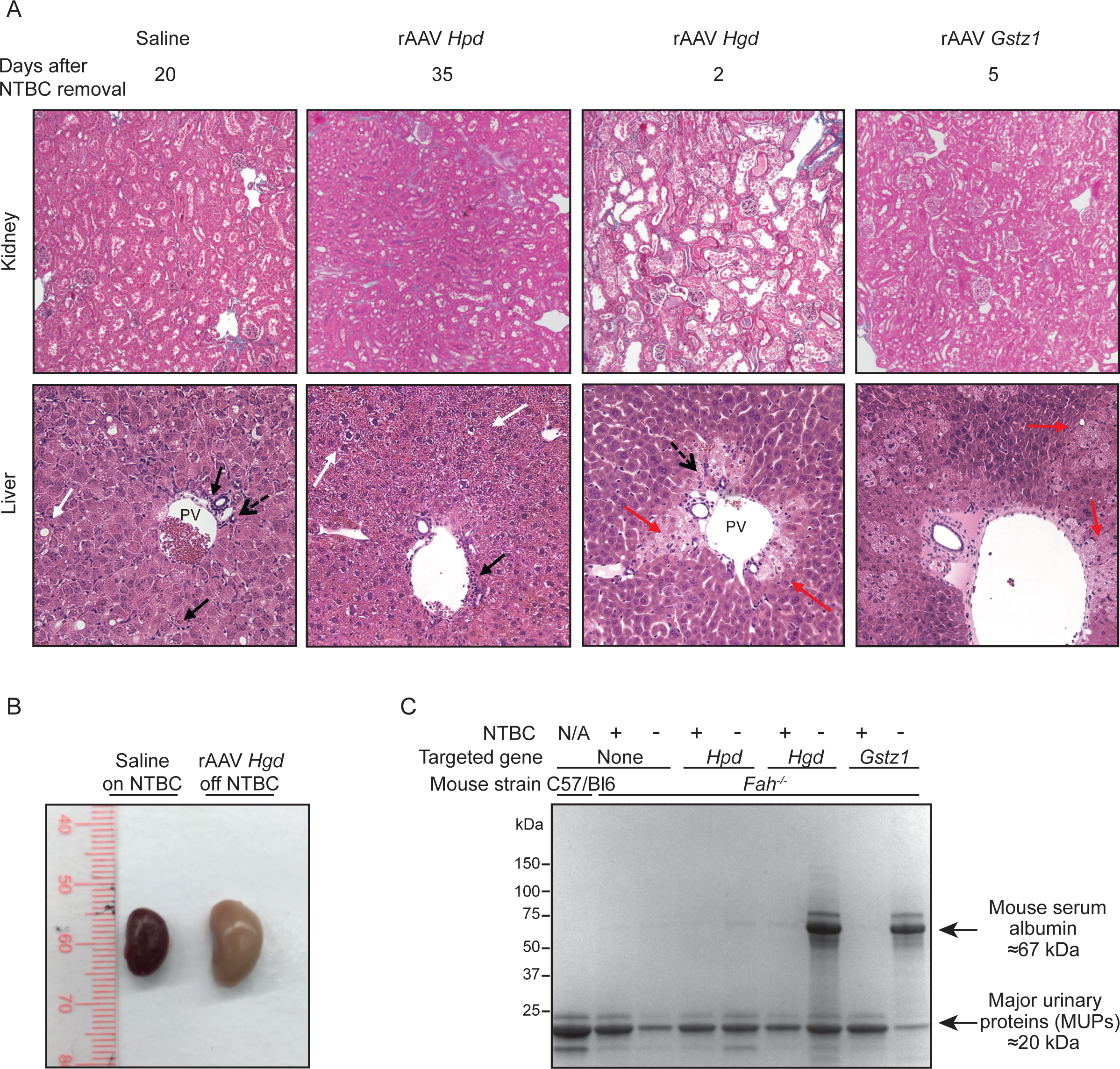
Tissue analysis of *Fah^-/-^* mice following metabolic rewiring by rAAV8-SaCas9. (A) Kidney and liver sections from *Fah^-/-^* mice treated with saline or rAAV8-SaCas9 vectors targeting *Hpd*, *Hgd,* or *Gstz1* as in Figure 2. *Fah^-/-^* mice injected with saline and the *Hpd, Hgd,* and *Gstz1* rAAV8s were sacrificed 20, 35, 2, and 5 days after NTBC removal, respectively. **Top panel**: Representative Masson’s trichrome-stained kidney sections from the treatment groups. Magnification: 10×. **Bottom panel:** Representative hematoxylin and eosin-stained liver sections from the treatment groups. PV: portal vein; black arrow: portal and lobular inflammation; dashed arrow: ductular proliferation; white arrow: necrosis; red arrow: ballooning degeneration. Magnification: 200×. (B) Representative kidneys from *Fah^-/-^*mice that were treated with saline and kept on NTBC (left) or treated with a vector targeting *Hgd* with NTBC withdrawn (right). (C) Urine samples (1µL per well) from treated *Fah^-/-^* animals on (+) and off (-) NTBC and untreated controls (as described in Figure 2) were loaded on a 4-15% gradient mini-PROTEAN TGX Stain-Free gel before electrophoresis, Coomassie staining, and imaging. Urine from a C57/Bl6 male mouse was used as a negative control. A band of the size expected for mouse serum albumin is indicated with an arrow. Also indicated with an arrow are bands of the expected size for the mouse major urinary proteins (MUPs).

Liver histology revealed that mice injected with the *Hgd*-targeting vector displayed substantial hepatocyte death in zones 1 and 2 with signs of apoptosis, moderate bile ductular proliferation, and portal inflammation. Hepatic steatosis and fibrosis were not detected (**Figure 3A, bottom panel)**. In mice injected with the *Gstz1-*targeting vector, mild lobular inflammation and significant and diffuse ballooning degeneration indicative of apoptotic hepatocyte death was observed (**Figure 3A, bottom panel)**. In animals injected with the *Hpd-*targeting vector, liver sections displayed moderate portal inflammation, mild ductular proliferation, and mild ballooning degeneration in zones 2 and 3, suggesting that gene-edited hepatocytes had not completely repopulated the diseased liver at the time of necropsy (**Figure 3A, bottom panel)**. Finally, liver histology revealed mild portal and lobular inflammation, mild ductular proliferation, glycogenated nuclei, and mixed steatosis in zone 2 in saline-injected controls (**Figure 3A, bottom panel)**. Collectively, these data indicate that liver-based editing can cause systemic effects impacting kidney function.

### Metabolic rewiring also occurs in “wild-type” mice

To ensure that the severely aggravated phenotypes observed in *Hgd*- and *Gstz1*-targeted animals were specific to the *Fah^-/-^* mouse model, we injected C57/Bl6N neonates with the three vectors or saline. We did not observe any changes in survival or glycemia between vector-treated animals and saline controls (**Figure 4A,B**). However, animals treated with the *Gstz1*-targeting vector had elevated urine SA levels, and animals injected with the *Hgd*- targeting vector had detectable HGA levels as expected from their respective metabolic blocks (**Figure 4C,D** and **Tables S5,S6**). Thus, this approach can partially recapitulate biochemical phenotypes associated with IEMs in wild-type animals, even when the metabolic block is limited to a single organ and not all hepatocytes are edited.

**Figure 4.**
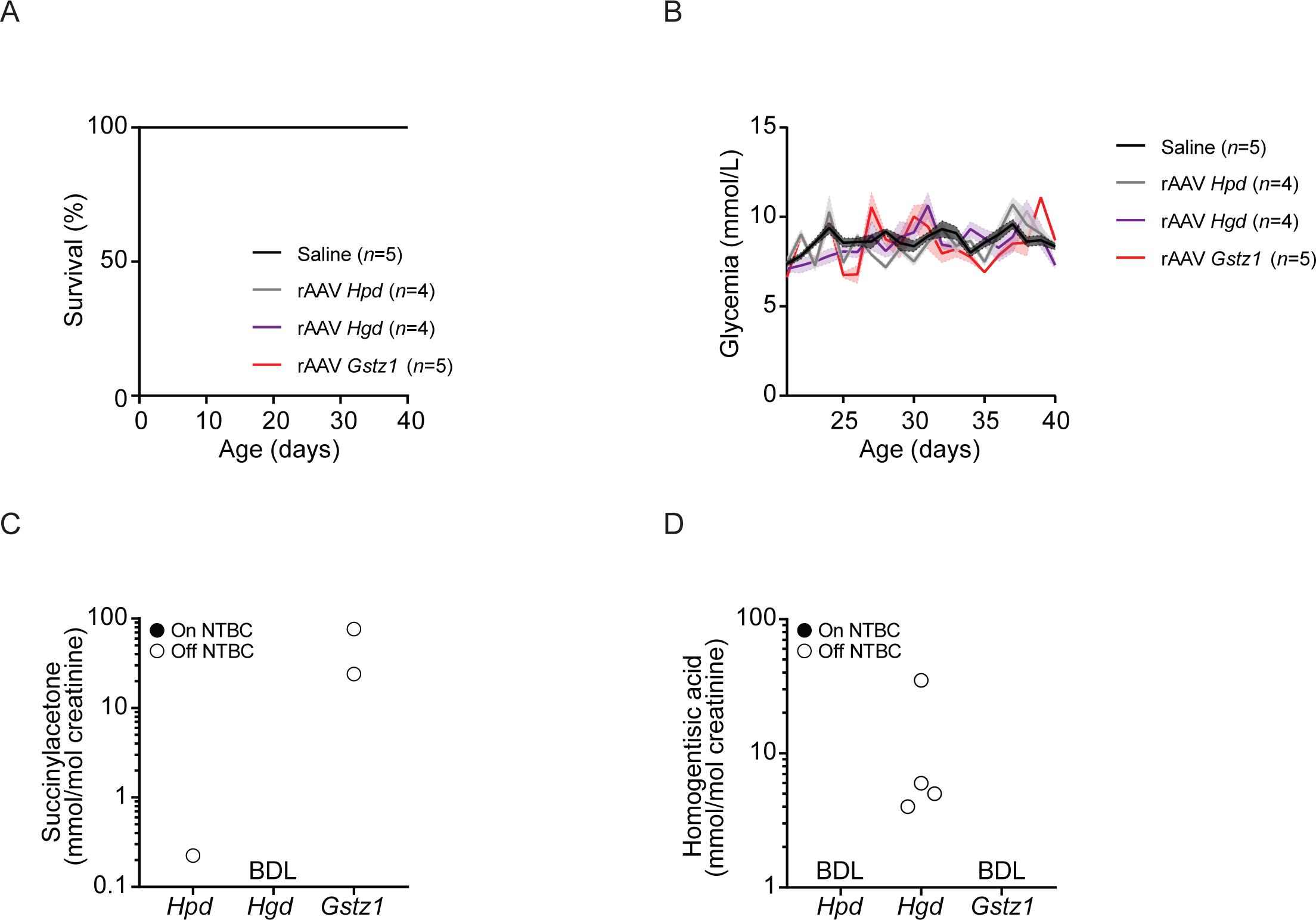
C57/Bl6N mice treated with the rAAV8-SaCas9 vectors have partial metabolic phenotypes and normal survival. (A) Survival analysis of male C57BL/6N mice injected at 2 days old with 5×10^10^ VGs rAAV8-SaCas9 or saline into the retro-orbital sinus. The numbers of mice per group (*n*) and rAAV8 targets are indicated. The animals treated with the vectors targeting *Hpd* and *Gstz1* are the same animals as in Figure 1C, whereas mice treated with the vector against *Hgd* are a separate group of animals injected solely for phenotypic studies. (B–D) Mice were assayed for phenotypic and metabolic modifications after weaning. (B) Glycemia of non-fasted mice. Solid lines indicate the mean and the shaded areas denote the SEM. Urine succinylacetone (C) and homogentisic acid (D) levels in mice treated as in (A) were determined 35 days after weaning. Samples were collected over 24 hours using metabolic cages containing 2-3 mice (**Tables S5 and S6**). The detection limits for succinylacetone and homogentisic acid were 0.1 mmol/mol and approximately 3 mmol/mol creatinine, respectively. BDL: below detection limit.

## DISCUSSION

Coupling rAAV-mediated *in vivo* gene delivery with CRISPR-Cas9 systems is a robust and rapid method to study gene functions in the somatic tissues of mice^45, 54, 61–63^. Here, we demonstrate that *in vivo* genome editing can modulate IEM-associated pathways to yield distinct phenotypes from those observed in whole-body knockout models. A limitation of this methodology is that gene disruption in somatic tissues is heterogeneous and incomplete, leaving a fraction of cells with altered functions that can affect the phenotype.

In *Fah^-/-^* mice, metabolic rewiring via *in vivo* editing recapitulated several hallmarks of HT-I. First, targeting *Hpd* rescued the lethal phenotype, as previously observed ^37, 39–41^. This rescue was maintained for at least 1 year post-NTBC removal. An increase in the editing level occurred over time in these mice likely due to the potent growth advantage of corrected hepatocytes following NTBC removal^32, 39, 41^. We also observed that liver-specific targeting of *Gstz1* in *Fah^-/-^* mice resulted in pronounced SA excretion and rapid death as in conventional double mutant *Fah*^-/-^ *Gstz1*^-/-^ mice^30^ (**Figure 5**). Interestingly, targeting *Hgd* resulted in the opposite phenotype compared to that observed in a classical knockout mouse model and when an FAH inhibitor was used to select *Hgd^-/-^* hepatocytes transplanted into wild-type recipient mice^38, 64^. While whole-body *Fah^-/-^ Hgd^-/-^* mice were protected from liver and renal damage, liver-specific inactivation of *Hgd* via *in vivo* editing in *Fah^-/-^* mice resulted in rapid death likely caused by kidney failure. This apparent discrepancy could be attributed to the fact that liver-specific *Hgd* disruption followed by NTBC withdrawal resulted in the rapid production of a massive amount of HGA, which is known to cause renal damage to *Fah^-/-^* mice^59, 65^. This HGA can be processed in the kidneys by active HGD and MAAI leading to the local accumulation of FAA and SA since FAH is not present in this organ in the *Fah^-/-^* mice (**Figure 5**). Previous work had shown that phenotypic rescue of HT-I in mice can occur by *Hgd* inactivation, an *in vivo* suppressor mutation^38^. In these spontaneous revertants, liver-function tests were normal but kidneys were pale and enlarged and showed extensive tubular damage^38^. In this original work, these data were not shown but they are reminiscent of our observations. It appears that the main difference is that our system rapidly produced a higher fraction of hepatocytes inactivated for *Hgd*, that created a bolus of HGA following sudden NTBC removal, which prevented any adaptation and caused acute renal failure (**Figure 5**). In HT-I, there is evidence that negative feedback loops can inhibit or down-regulate upstream enzymes in the tyrosine degradation pathway. For example, in *Fah^-/-^* mice, TAT mRNA levels are markedly reduced without NTBC treatment^23^ (**Figure 1A**). In humans, HPD’s enzymatic activity is greatly reduced in patients with HT-I compared to healthy controls^17, 66^. These compensatory mechanisms may partially protect liver and kidney cells from the toxic accumulation of FAA and SA and may have been bypassed by the sudden perturbations created by the editing process and NTBC withdrawal.

**Figure 5.**
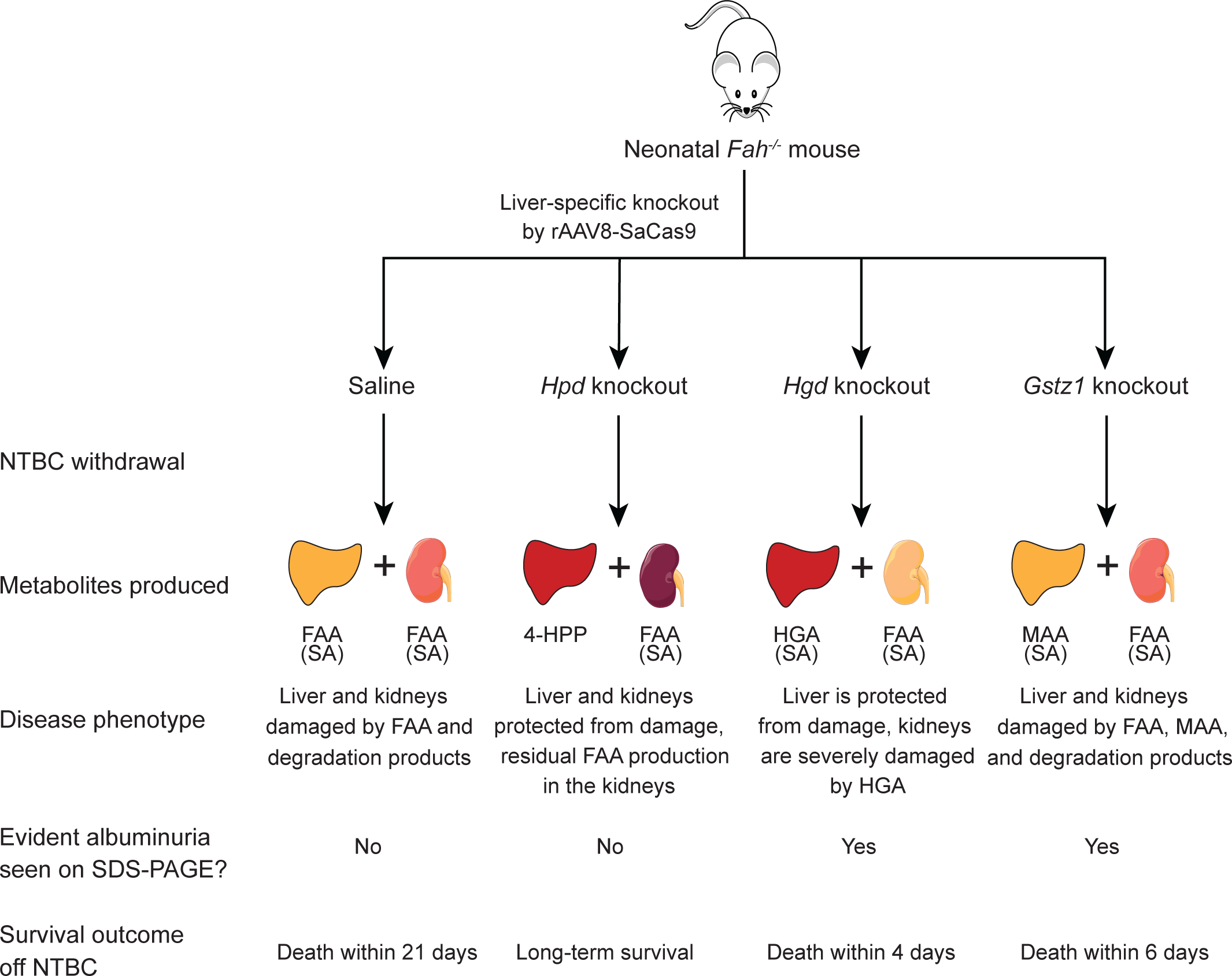
Model of metabolic rewiring in *Fah^-/-^* mice following liver-specific knockout of tyrosine catabolic enzymes. Observed disease phenotypes and survival outcomes after injecting neonatal *Fah^-/-^* mice with saline or liver-specific rAAV8-SaCas9 vectors targeting *Hpd*, *Hgd*, or *Gstz1* and NTBC withdrawal. The main metabolites predicted to be produced by the liver and unedited kidneys in each condition are shown. Fumarylacetoacetate (FAA) and maleylacetoacetate (MAA) accumulation leads to non-enzymatic production of succinylacetone (SA) (see Figure 1A).

We propose a model of metabolic interplay between the edited liver and the non-edited kidneys in *Fah^-/-^* mice following NTBC withdrawal (**Figure 5**). Of note, this model does not exclude the possibility that circulating HGA can also re-enter non-targeted, HGD- expressing hepatocytes, and cause their rapid death^67, 68^. Irrespective of the physiological mechanism, the differences between our observations and those of whole-body knockout models highlight the importance of using tissue-specific genome editing in animal models of human genetic disorders to investigate its systemic impacts. It has recently been shown that metabolic pathway rewiring following liver-directed CRISPR-Cas9 knockout can be used to rescue glutaric acidemia type 1, an inborn error of metabolism affecting the lysine degradation pathway^69^. *In vivo* genome editing is thus a promising approach that could be used to dissect metabolic pathways affected in other IEMs.

Other than the limitation imposed by the impossibility to edit all hepatocytes within the liver, even in mice, there may be major differences in phenotypes between humans and mice deficient in the same gene product. A prime example is that HT-I mice have a neonatal lethal phenotype and are not tyrosinemic (i.e. they do not have elevations of plasma tyrosine)^27^, unless treated with NTBC^23^, contrary to humans with tyrosinemia type I. Our studies have also been performed in neonates which may have differences in liver functions compared to adult mice. Finally, for historical reasons^52, 53, 70^, we only used male mice in this study even though sexual dimorphism in response to similar genetic treatments had been reported^71–73^. Reassuringly, we have observed that *in vivo* genome editing using *Streptococcus thermophilus* CRISPR1-Cas9 (St1Cas9) shows no such bias when targeting *Hpd* in this same mouse model^41^.

## METHODS

### Genome editing vectors

The CMV-driven SaCas9 nuclease vector pX601^43^ (Addgene plasmid #61591) was a gift from Feng Zhang (Massachusetts Institute of Technology). Target sequences for *Hpd*, *Hgd*, and *Gstz1* were designed using the web-based CRISPR design tool CRISPOR^48^. The sgRNA sequences used are listed in **Table S1**. When required, the sgRNA sequence was modified to encode a G at position 1, to meet the transcription initiation requirement of the human U6 promoter. Following *in vitro* screening, selected SaCas9 sgRNAs were cloned into the TBG-driven SaCas9 nuclease rAAV vector pX602^43^ (Addgene plasmid #61593; also a gift from Feng Zhang) for *in vivo* gene editing. The inverted terminal repeat integrity of the rAAV vector pX602 was assessed by BssHII digestion.

### Cell culture and transfection

Neuro2A cells were obtained from the ATCC (CCL-131, Manassas, VA, USA) and maintained at 37°C under 5% CO2 in Dulbecco’s modified Eagle’s medium (DMEM, high glucose, GlutaMAX™ Supplement) (Thermo Fisher Scientific, Waltham, MA, USA) supplemented with 10% fetal bovine serum (FBS, Gibco, Thermo Fisher Scientific, Waltham, MA, USA), 1% penicillin-streptomycin (Thermo Fisher Scientific, Waltham, MA, USA). The cells were tested and found negative for mycoplasma contamination. Cells (2×10^5^) were transfected with 500 ng of pX601 (expressing both the sgRNA and SaCas9) using an Amaxa 4D-Nucleofector (Lonza, Basel, Switzerland) according to the manufacturer’s recommendations and harvested 3 days post-transfection.

### Surveyor and TIDE assays

Genomic DNA was extracted from 2.5×10^5^ Neuro2A cells with 250 μL of QuickExtract DNA Extraction Solution (Lucigen, Middleton, WI, USA) or from 30 mg of mouse liver using a EZ-10 Spin Column Animal Genomic DNA Miniprep Kit (Bio Basic, Markham, ON, CA), per the manufacturers’ recommendations. Loci were amplified by polymerase chain reaction (PCR) using the primers listed in **Table S7**. Surveyor assays were performed with the Surveyor Mutation Detection Kit (Integrated DNA Technologies, Coralville, IA, USA) as described^55^. Samples were resolved on 10% polyacrylamide gels in Tris-borate-EDTA buffer and bands were visualized with RedSafe Nucleic Acid Staining Solution (iNtRON Biotechnology, Seongnam, South Korea). Gels were imaged using a ChemiDoc MP Imaging System (Bio-Rad, Hercules, CA, USA) and bands were quantified using Image Lab Software (Bio-Rad). TIDE analysis was performed using a significance threshold value for decomposition of *P* < 0.001^57^.

### Adeno-associated virus production

The rAAV8s were produced by the Canadian Neurophotonics Platform’s Viral Vector Core (The Molecular Tools Platform) using the triple plasmid transfection method, as described^74^. Briefly, HEK293T17 (ATCC CRL-11268, Manassas, VA, USA) cells were transfected using polyethylenimine (Polysciences) with the helper plasmid pxx-680 (a gift from R. Jude Samulski, University of North Carolina), the rep/cap hybrid plasmid pAAV2/8 (Addgene #112864, a gift from James Wilson, University of Pennsylvania), and the rAAV vector pX602. After 24 hours, the medium was replaced with medium without FBS, and the cells were harvested 24 hours later. We purified rAAV particles from the cell extracts using freeze/thaw lysis followed by a discontinuous iodixanol gradient. Viruses were resuspended in phosphate-buffered saline containing 320 mM NaCl, 5% D-sorbitol, and 0.001% pluronic acid (F-68), aliquoted, and stored at -80°C. The rAAVs were titrated by quantitative PCR using LightCycler 480 SYBR Green I Master mix (Roche, Basel, Switzerland) and inverted terminal repeat primers as described^75^. Vector yields were 1×10^13^–3×10^13^ VG/mL. The purity of viral preparations was determined by sodium dodecyl sulfate (SDS)-polyacrylamide gel electrophoresis on a 4– 15% Mini-PROTEAN TGX Stain-Free Gel (Bio-Rad, Hercules, CA, USA) in Tris-glycine-SDS buffer (**Figure S3**).

### Animal experiments

*Fah^-/-^* mice^27^ in a C57BL/6 genetic background were a kind gift from Robert Tanguay (Laval University). C57BL/6N mice were purchased from Charles River (Strain code 027) (Wilmington, MA, USA). All mice were group-housed and fed a standard chow diet (Harlan #2018SX) with free access to food and water. For *Fah^-/-^* mice, the drinking water was supplemented with 7.5 mg/L NTBC. Mice were exposed to a 12:12-h dark-light cycle and kept at an ambient temperature of 23±1°C. Animals were cared for and handled according to the *Canadian Guide for the Care and Use of Laboratory Animals.* The Laval University Animal Care and Use Committee approved the procedures.

Neonatal (2-day-old) male mice were injected intravenously in the retro-orbital sinus^76^ with saline or 5×10^10^ VGs of an rAAV8, adjusted to an injection volume of 20 µL with saline. *Fah^-/-^* mice were weaned at 21 days old and NTBC was removed at the indicated time points. Body weight and glycemia were monitored post-NTBC removal. Glycemia was measured in unfasted mice between 9 and 10 am. Animals were sacrificed by cardiac puncture under anesthesia at predetermined time points or when they had lost 20% of their body weight. The majority of the liver and one kidney were snap frozen, while a portion of the liver and the other kidney was fixed in 4% paraformaldehyde.

### Urine collection and SDS-PAGE analysis

Urine was collected from groups of 2–5 mice overnight using metabolic cages (Tecniplast) before and at different time points after NTBC removal. Urine was centrifuged at 900 x *g* for 5 minutes, aliquoted, and frozen at -80°C for downstream applications. For SDS-PAGE analysis, 5µL of urine per mouse was loaded on a 4–15% Mini-PROTEAN TGX Stain-Free Gel (Bio-Rad, Hercules, CA, USA) before Coomassie staining with QC Colloidal Coomassie (Bio-Rad) and imaging using a ChemiDoc MP Imaging System (Bio-Rad).

### Histology

Liver portions and kidneys were fixed in 4% paraformaldehyde for 24 hours, then dehydrated in 70% ethanol, embedded in paraffin, and sliced into 4 µm sections, which were mounted on slides. Liver sections were stained with hematoxylin and eosin and reviewed by a blinded pathologist and a blinded hepatologist at a magnification of 200X. Kidney sections were stained with Masson’s trichrome and reviewed by a blinded nephrologist using an Olympus BX45 microscope at different magnifications.

### SA and HGA quantification

Urine SA and HGA levels were quantified by gas chromatography-mass spectrometry (GC-MS), as previously described^77, 78^. All analyses were performed in the biochemical genetics laboratory at the CHUS.

## DECLARATIONS

### Ethics approval

Animals were cared for and handled according to the *Canadian Guide for the Care and Use of Laboratory Animals.* The Laval University Animal Care and Use Committee approved the procedures.

### Availability of data and materials

The datasets used and analyzed during the current study are available from the corresponding author upon request.

### Author’s contributions

Conceptualization, J-F. R., S.C., P.J.W., M.P., and Y.D.; Methodology, J-F. R., S.C., D.C., P.J.W., D.D., F.M., M.P., and Y.D.; *In vitro* and *in vivo* genome editing, mouse experiments, data and sample collection: J-F. R., S.C., and C.G.; Kidney histology: R.V.U., F.M.; Liver histology: D.D., M.P.; SA and HGA quantification: D.C., P.J.W.; Writing – Original Draft, J-F. R., S.C., P.J.W., and Y.D.; Writing – Review and Editing, J-F. R., S.C., F.M., P.J.W., M.P., and Y.D.; Supervision, F.M., P.J.W., M.P., and Y.D.; Funding Acquisition, Y.D. All authors read and approved the final manuscript.

## Acknowledgments

We are indebted to Robert Tanguay for providing the HT-I mouse model, NTBC, and his expertise and support. We also thank Marie-Ève Paquet and the Viral Vector Core staff at the Canadian Neurophotonics Platform for rAAV8 production. We thank Patrick Bherer, Tommy Gagnon, and the technologists of the CHUS biochemical genetics laboratory for help with biochemical assays, and High-Fidelity Science Communications for editing the manuscript.

