## Supplemental Figures and Tables for "*In vivo* dissection of the mouse tyrosine catabolic pathway with CRISPR-Cas9 identifies modifier genes affecting hereditary tyrosinemia type 1"

Figure S1 | Screening for active SaCas9 sgRNAs in mouse Neuro-2a cells and TIDE assays in treated *Fah*$^{-/-}$ mice

Figure S2 | Long-term follow-up of *Hpd*-targeted mice after removing NTBC

Figure S3 | Purified rAAV8 vectors used in this study

**SUPPLEMENTAL TABLES**

Table S1 | SaCas9 guide RNA (spacer) sequences

Table S2 | Raw urine succinylacetone quantification data from *Hpd*-targeted animals 1 year after NTBC removal

Table S3 | Raw urine succinylacetone quantification data from *Fah*$^{-/-}$mice

Table S4 | Raw urine homogentisic acid quantification data from *Fah*$^{-/-}$ mice

Table S5 | Raw urine succinylacetone quantification data from C57/Bl6N mice

Table S6 | Raw urine homogentisic acid quantification data from C57/Bl6N mice

Table S7 | PCR primers used in Surveyor and TIDE assays and their amplicon sizes

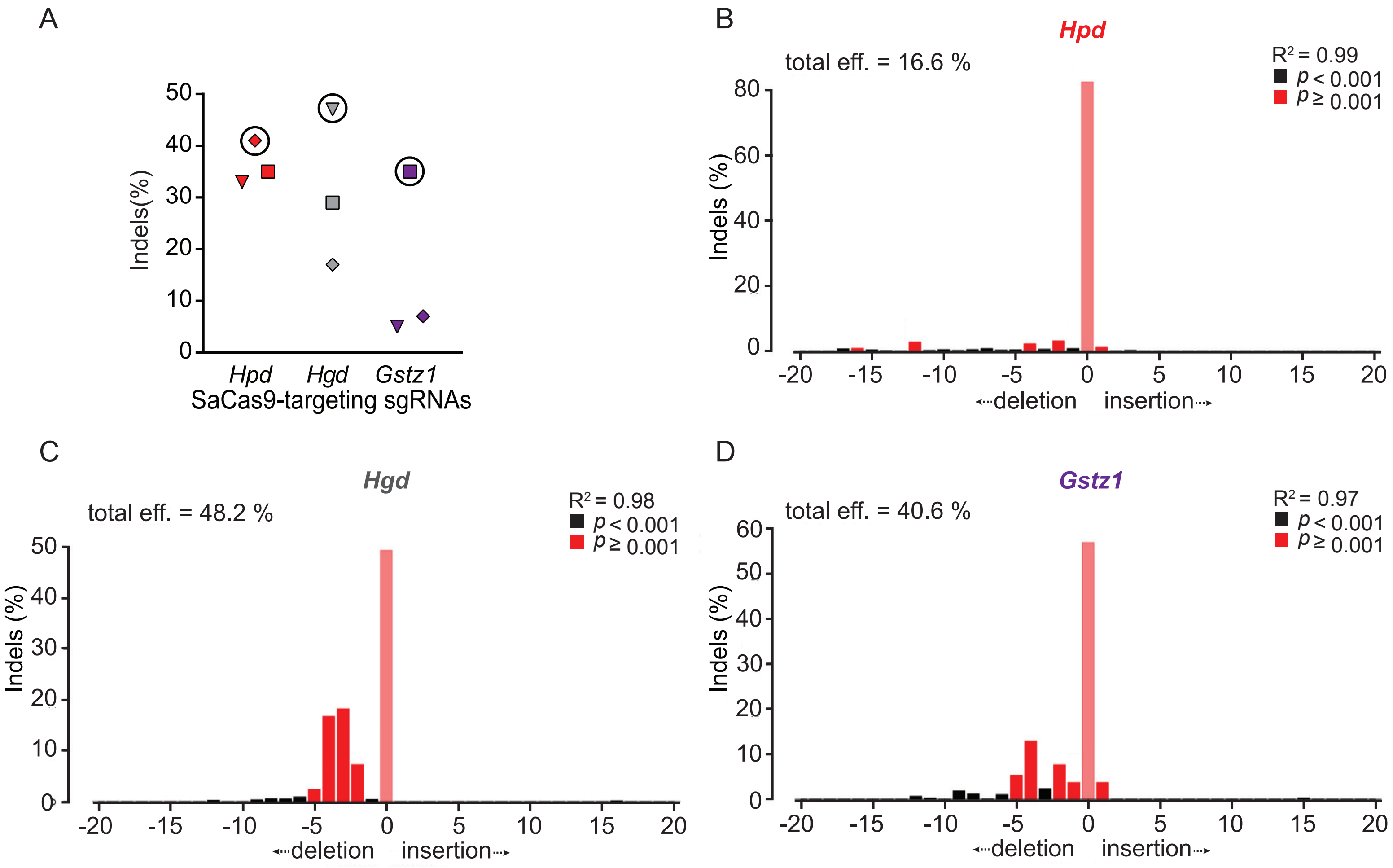


**Rivest et al. Figure S1**

**Figure S1 | Screening for active SaCas9 sgRNAs in mouse Neuro-2a cells and representative *in vivo* indels profiles** (A) Surveyor assays to determine the activities of sgRNAs targeting SaCas9 to *Hpd, Hgd*, and *Gstz1*. Neuro-2a cells were transiently transfected with 500ng of a single vector expressing SaCas9 and each sgRNA. Each symbol represents a different sgRNA. Surveyor assays were performed 3 days later to determine the frequency of SaCas9-induced insertions and deletions (indels). A vector encoding EGFP was used as a negative control (not shown). Circles indicate the sgRNAs chosen for *in vivo* studies. (B-D) Representative TIDE analyses of *Fah*$^{-/-}$ mice injected with rAAV8-SaCas9 vectors targeting *Hpd* (B), *Hgd* (C), and *Gstz1* (D) as described in **Figure 1D**.

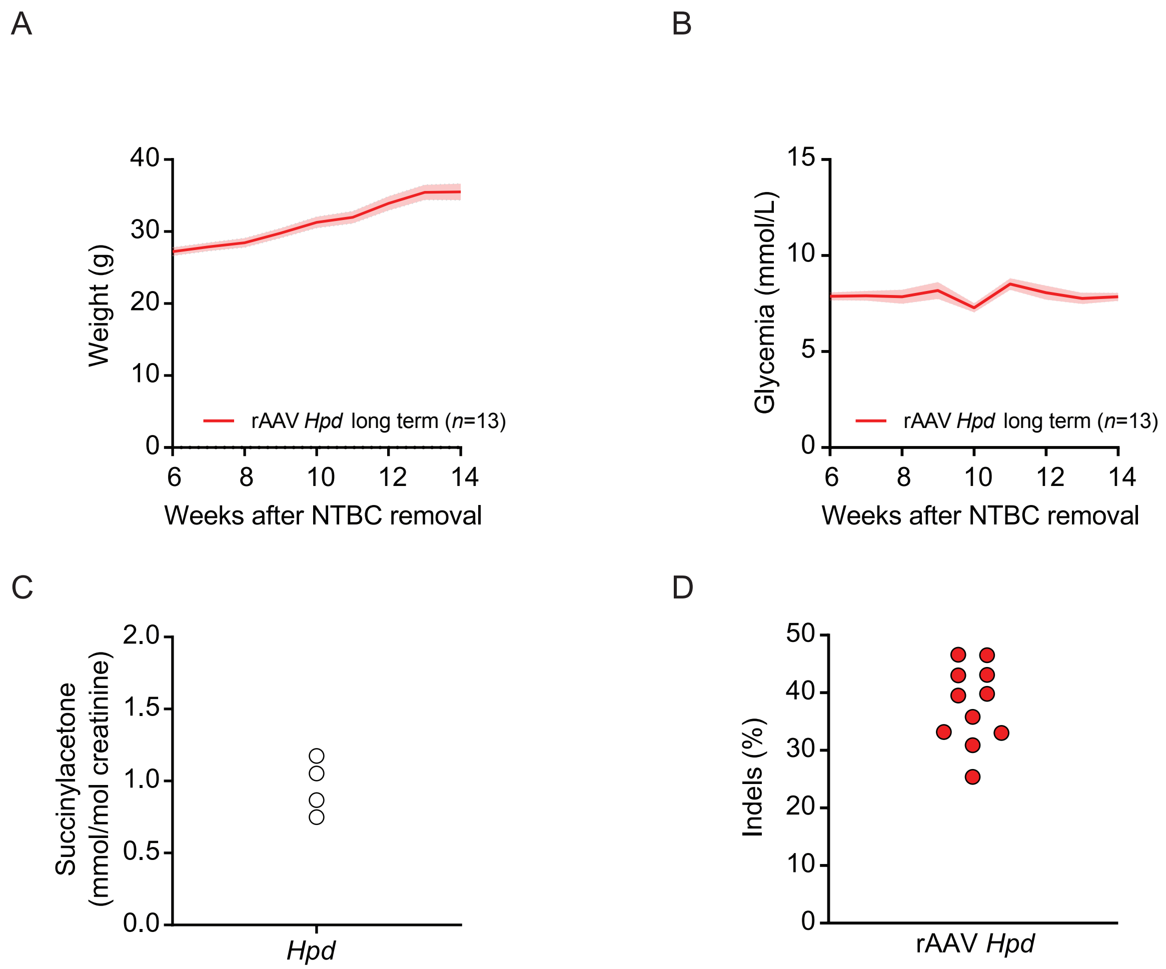


Rivest et al. Figure S2

**Figure S2 | Long-term follow-up of *Hpd*-targeted mice after removing NTBC** (A) Long-term weights and (B) glycemia of the mice taken off NTBC in **Figure 2**, from weeks 6–14 post-removal. (C) One year after NTBC removal, urine samples were collected over 24 hours using metabolic cages and succinylacetone levels were measured. The detection limit was 0.1 mmol/mol creatinine. Raw data are presented in **Table S2**. (D) After sacrifice, genomic DNA was extracted from the livers of each mouse and TIDE assays were used to determine the indel frequency. Each symbol represents a different animal. A mouse injected with saline was used as a negative control for the TIDE assay.

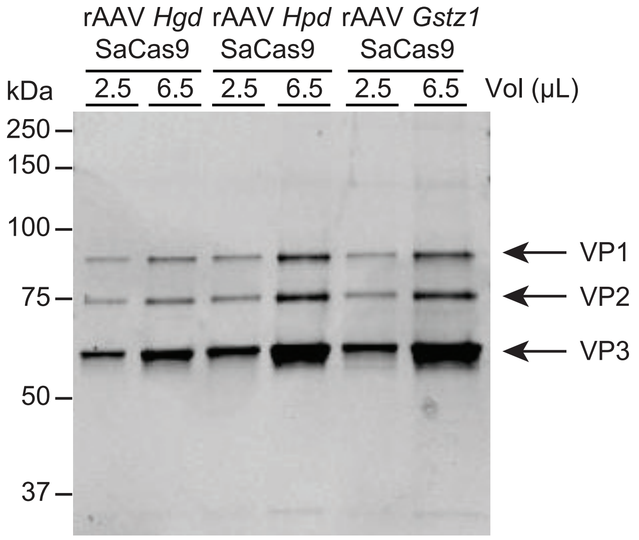


**Rivest et al. Figure S3**

**Figure S3 | Purified rAAV8 vectors used in this study** (A) Fixed volumes of the purified rAAV8-SaCas9 vectors used in the study were resolved on mini-PROTEAN TGX Stain-Free Gels. The molecular weights (kDa) of protein standards are indicated. The three rAAV capsid proteins VP1, VP2 and VP3 are indicated by an arrow.

**Table S1 | SaCas9 guide RNA (spacer) sequences**

| Gene | Exon | Target size (bp) | PAM | Target | Indels in the Surveyor assay (%) |
|---|---|---|---|---|---|
| *Hpd* | 3 | 20 | CCGAGT | **G**ATTGCCAACCCAGAAGGTCA | 35 |
| | 7 | 20 | GTGAGT | **GAGTTTGCTGTGCTGCAGACG** | **41** |
| | 10 | 20 | CGGAGT | GGACGACACGCAGGTGCACA | 33 |
| *Hgd* | 9 | 20 | TTGAGT | **GGAGGTCTATGGTGTCCACT** | **47** |
| | 12 | 21 | GGGAGT | **G**TGTCATCTTCCCACCTCGGTG | 29 |
| | 13 | 21 | GTGGAT | GTCACTATGAGGCAAAGCAAG | 17 |
| *Gstz1* | 4 | 20 | GTGAGT | **GAATCACCATTGTCCAGTCA** | **35** |
| | 5 | 20 | GTGGGT | GCGCACGATGGCTCTTTTCT | **5** |
| | 8 | 20 | ATGGAT | GTACTAAGCACACATCAGCC | **7** |

Target sequences for *Hpd*, *Hgd* and *Gstz1* were selected using CRISPOR. We selected sgRNAs with high predicted activity and low predicted off-target scores. When required, the sgRNA sequence at position 1 was changed to 'G' to meet the transcription initiation requirements of the human U6 promoter (indicated in bold red). The target sequences in bold were chosen for *in vivo* studies. Underlined blue nucleotides indicate mismatches with the human genome sequence for one of the chosen targets.

**Table S2 | Raw data from urine succinylacetone quantification in *Hpd*-targeted animals 1 year after NTBC removal**

| Group | Units | Cage 1 | Cage 2 | Cage 3 | Cage 4 |
|---|---|---|---|---|---|
| *Hpd* long-term | mmol/mol creatinine (number of animals per cage) | 1.17 (3) | 1.05 (3) | 0.75 (4) | 0.87 (3) |

Urine was collected overnight before and at different time points after NTBC removal from groups of 3–4 mice as described in **Figure S2**. Succinylacetone levels were quantified by GC-MS.

**Table S3 | Raw urine succinylacetone quantification data from *Fah*$^{-/-}$ mice**

| Group | Units | On NTBC | | | | Off NTBC | | | |
|---|---|---|---|---|---|---|---|---|---|
| | | Cage 1 | Cage 2 | Cage 3 | Cage 4 | Cage 1 | Cage 2 | Cage 3 | Cage 4 |
| Saline | mmol/mol creatinine (number of animals per cage) | 8.05 (4) | 20.18 (3) | 9.59 (3) | 9.24 (5) | 74.77 (3) | 53.16 (2) | | |
| *Hpd* | | 5.49 (4) | 7.58 (5) | 7.09 (3) | | 0.51 (3) | 0.51 (4) | 0.50 (3) | 0.52 (4) |
| *Hgd* | | 5.40 (5) | 5.47 (4) | 12.17 (2) | | 24.15 (3) | 16.77 (5) | | |
| *Gstz1* | | 10.06 (3) | 11.97 (3) | 11.18 (5) | | 82.23 (3) | 133.35 (3) | 117.28 (2) | |

Urine was collected overnight before and at different time points after NTBC removal from groups of 2–5 mice as described in **Figure 2**. Succinylacetone levels were quantified by GC-MS.

**Table S4 | Raw urine homogentisic acid quantification data from *Fah*$^{-/-}$ mice**

| Group | Units | On NTBC | | | Off NTBC | | |
|---|---|---|---|---|---|---|---|
| | | Cage 1 | Cage 2 | Cage 3 | Cage 1 | Cage 2 | Cage 3 |
| Saline | mmol/mol creatinine (number of animals per cage) | BDL (4) | BDL (3) | | BDL (3) | BDL (2) | |
| *Hpd* | | BDL (4) | BDL (5) | BDL (3) | BDL (3) | BDL (4) | BDL (3) |
| *Hgd* | | 22 (5) | 4 (4) | | 1613 (3) | 1440 (5) | |
| *Gstz1* | | BDL (3) | BDL (3) | BDL (5) | BDL (3) | BDL (3) | BDL (2) |

Urine was collected overnight before and at different time points after NTBC removal from groups of 2–5 mice as described in **Figure 2.** Homogentisic acid levels were quantified by GC-MS.
BDL: Below detection limit

**Table S5 | Raw urine succinylacetone quantification data from C57/Bl6N mice**

| Group | Units | Cage 1 | Cage 2 |
|---|---|---|---|
| *Hpd* | mmol/mol creatinine (number of animals per cage) | 0.23 (2) | |
| *Hgd* | | BDL (3) | BDL (3) |
| *Gstz1* | | 76.67 (2) | 24.07 (2) |

Urine was collected overnight before and at different time points after NTBC removal from groups of 2–3 mice as described in **Figure 4**. Succinylacetone levels were quantified by GC-MS.
BDL: Below detection limit

**Table S6 | Raw data from urine homogentisic acid quantification in C57/Bl6N mice**

| Groups | Units | Cage 1 | Cage 2 | Cage 3 | Cage 4 |
|---|---|---|---|---|---|
| *Hpd* | mmol/mol creatinine (number of animals per cage) | BDL (2) | | | |
| *Hgd* | | 5 (3) | 35 (3) | 6 (2) | 4 (2) |
| *Gstz1* | | BDL (2) | | | |

Urine was collected overnight before and at different time points after NTBC removal from groups of 2–3 mice as described in **Figure 4**. Homogentisic acid levels were quantified by GC-MS.
BDL: Below detection limit

**Table S7 | PCR primers used in Surveyor and TIDE assays and their amplicon sizes**

| Target | Primer | Size (bp) |
|---|---|---|
| *Hpd* exon 3 Forward | GTCACCCATACTGTTCTCACGTA | 466 |
| *Hpd* exon 3 Reverse | CAAGGTTCCAAAGTGCCAGTCC | |
| *Hpd* exon 7 Forward | GCAGGCGCAGTGCCCAAGACAC | 498 |
| *Hpd* exon 7 Reverse | CAGCACATGCCCAGGTCACATGG | |
| *Hpd* exon 10 Forward | GTGTAACGGGTGTATGCTCAATG | 452 |
| *Hpd* exon 10 Reverse | GTGATGATGTCTTCCGTCTTGAG | |
| *Hgd* exon 9 Forward | TGAGTTGTGGCTAACTGGGG | 420 |
| *Hgd* exon 9 Reverse | AGGCAGGCATTTTGTCTAAGGA | |
| *Hgd* exon 12 Forward | CTGTCACTTGAAAGCACCCT | 451 |
| *Hgd* exon 12 Reverse | TCACACTCTCCCAGCCTGTC | |
| *Hgd* exon 13 Forward | GTGCTATTGTGGAATAGTTG | 412 |
| *Hgd* exon 13 Reverse | TCCTCACTCCACCTCTGTGA | |
| *Gstz1* exon 4 Forward | GACCACAGTAAAGAGTACAGGGA | 400 |
| *Gstz1* exon 4 Reverse | GCTTGGTCACTTGTAGGTTAGTT | |
| *Gstz1* exon 5 Forward | GGAGTTTGCTGCCTCTCCCCTC | 505 |
| *Gstz1* exon 5 Reverse | GCTACAGAGCAGATGACCAGGAG | |
| *Gstz1* exon 8 Forward | GGGAAACTTGACATGGGAGAAAT | 393 |
| *Gstz1* exon 8 Reverse | CAGTTGATAATGGCCTGGTGTAG | |
